# Attenuation of mosaic aneuploidy and erroneous first mitotic division of human embryos

**DOI:** 10.1101/2025.04.05.647367

**Authors:** Cerys E. Currie, Aleksandra Byrska, Deborah M. Taylor, Muriel Erent, Daniela Bakalova, Xuhui Sun, Constandina Koki, Nigel J. Burroughs, Richard. A. Anderson, Adele L. Marston, Geraldine M. Hartshorne, Andrew D. McAinsh

## Abstract

Human reproduction is inherently inefficient^1^ and 1 in 6 people worldwide suffer infertility. In vitro fertilisation (IVF) can help some couples conceive, but only ~30% of cycles are successful. One factor affecting IVF efficacy is mitotic-origin (mosaic) aneuploidy in which embryos contain a mixture of cells with different numbers of chromosomes^2^. We previously showed that chromosome segregation error phenotypes are frequent in the first mitotic division of the human embryo^3^. However, the cause of these errors and impact on daughter cell karyotype is unknown. Here, using live chromosome imaging and next generation sequencing we show that activation of the microtubule depolymerase KIF2C reduces chromosome segregation errors and mitotic-origin aneuploidy at the 2-cell stage. The number of first divisions that show alternative cleavage patterns (associated with failed embryo development in IVF clinics) are also reduced with KIF2C activation. Our findings demonstrate that modulation of microtubule dynamics is a potential therapeutic route to improving human embryo quality and IVF outcomes.

## Introduction

Infertility is generally defined as the failure to achieve pregnancy after 1 year of trying to conceive, and is estimated to affect 1 in 6 reproductive-aged individuals worldwide^4^.

Infertility and its associated treatment create a significant negative impact on society, causing emotional distress, economic hardship and pressure on health care systems. Pioneering in vitro fertilisation (IVF) approaches provided a route to treat infertility, although success rates have not increased above ~30%, and worsen with increasing maternal age. Combined with a cultural shift towards first pregnancies at later stages of life, the decline in global birth rate is leading to mounting economic pressure from an ageing demographic with looming social, economic and geopolitical consequences^5,6^.

It is widely accepted that only ~30% of human conceptions result in live birth^7^. There is a growing need to understand the molecular basis of IVF failure and design therapeutic strategies to increase success rates – currently none exist. It is well established that eggs and embryos having the wrong number or structure of chromosomes (aneuploidy) is a leading cause of reproductive failure^7,8^. There are two origins of numerical aneuploidy: 1) Chromosome segregation errors during the female meiotic divisions that generate the egg. Problems in meiosis increase with advancing maternal age, resulting in embryos in which all cells are aneuploid^9^. This is associated with failed embryo development and implantation, clinical miscarriage^8^ or developmental syndromes^10^. 2) Chromosome segregation errors in the post-fertilisation mitotic divisions. These lead to mosaic aneuploidy where different cells in an embryo contain different chromosome complements; some cells can be normal.

Mosaicism occurs independently of maternal age^11^ and meiotic aneuploidy^12^. Remarkably, three-quarters of human cleavage-stage embryos contain unbalanced aneuploidies of mitotic origin^13–15^, and single cell data shows that >80% of human blastocysts can be classed as mosaics, with ~20% cells containing aneuploidies^12,16^. Moreover, clinical trials have shown that blastocysts diagnosed by PGT-A as low or mid-level mosaic (20-50% aneuploid cells) are of identical developmental potential as those diagnosed as euploid^17^. Indeed, some embryos diagnosed euploid are in fact mosaics^12^, and the status of mosaic embryos remains an area of heavy debate in clinical practice^18^. However, catastrophic mitotic-origin aneuploidy, affecting multiple chromosomes, is non-viable and is associated with developmental arrest prior to blastocyst formation^19,20^.

Our previous work identified aneuploidy-generating phenotypes during the first mitotic division of human embryos^3^. These included multipolar chromosome segregation and lagging chromosomes that gave rise to micronuclei - likely origins of mosaic numerical and structural aneuploidy. The presence of lagging chromosomes is thought to be the result of chromosome mis-attachments and has been reported in other large mammalian embryos undergoing the first mitosis (*e*.*g*. cow^21^), but interestingly not mouse where embryonic aneuploidy is minimal^22^. Erroneous mitotic phenotypes have also been observed at the blastocyst stage in humans, suggesting chromosomal instability is a feature of cells within the human preimplantation embryo^23,24^.

The molecular mechanisms underlying the high rate of errors remains poorly understood, and experimentation is challenging due to the scarcity of early human embryos available for research and strict regulations on genetic manipulation. Some progress is, however, being made: human zygotes have been shown to have problems maintaining a bipolar spindle, and this is implicated in the formation of multinucleated daughter cells^25^. On the other hand, deregulated DNA replication and DNA damage during S-phase in the zygote is associated with mosaic structural aneuploidies and chromosome loss^26,27^. Neither study explains the origin of chromosome mis-attachments, and no other work has connected these potentially aneuploidy-generating phenotypes with karyotype change in resulting daughter cells.

### Activating the KIF2C microtubule depolymerase reduces error-generating phenotypes in the first mitotic division

Low-fidelity mitoses, featuring lagging chromosomes and multipolar divisions, are a common feature of cancer cells and human cells with increased developmental potential^28–30^. In these settings, errors in chromosome attachment to the mitotic spindle - resulting from hyperstable microtubule-kinetochore attachments - can cause segregation errors and aneuploidy^31^. We reasoned that a similar mechanism may be underlying the high error rate during the first mitotic division of the human zygote. To investigate this, we created human embryos for research with donated MII oocytes using intra-cytoplasmic sperm injection (ICSI), and obtained zygotes deselected from patient treatment. Embryos where fertilisation by multiple sperm was suspected (eg. 3PNs) were discarded from our dataset. Consistent with our previous findings, we observed chromosome segregation errors during the first mitosis (Fig. 1a, top panel). Next, we treated human embryos with UMK57, a small molecule agonist for the molecular motor KIF2C^28,31–33^. KIF2C (also known as MCAK) is a member of the kinesin-13 family that uses an ATP-dependent mechanism to remove tubulin subunits from the ends of microtubules and promote their depolymerisation^34^. Activation of KIF2C has been shown to increase turnover of microtubule-kinetochore attachments and reduce the frequency of lagging chromosomes in cancer, stem and aged human cells^35,36^.

**Figure 1.**
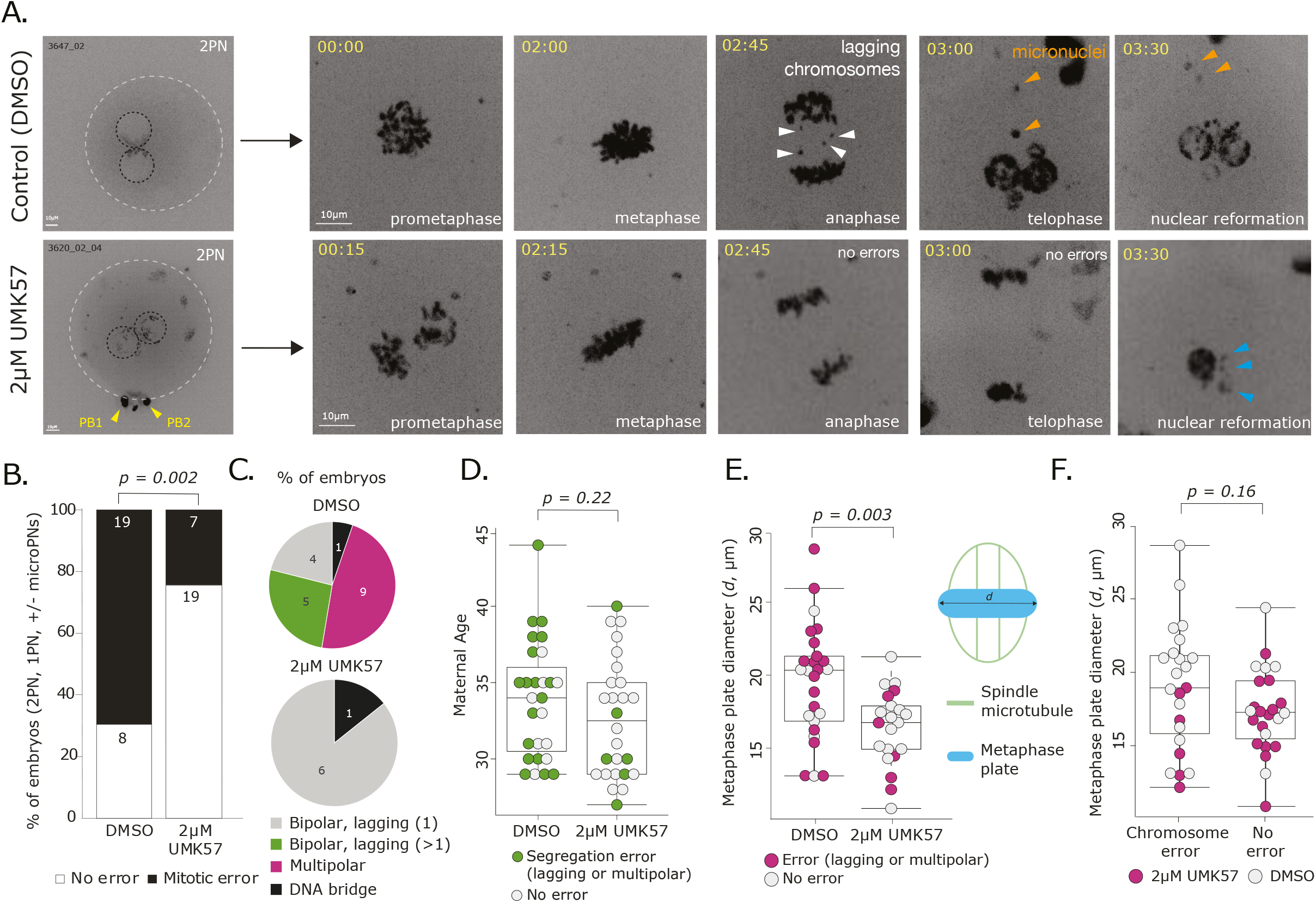
Rescue of mitotic errors in the first cleavage division of human embryos. a) Representative time lapse imaging of representative human embryos (DMSO: 3647_02, 2µM UMK57: 3620_02_02) progressing through the first embryonic mitosis. Chromosomes are visualised using SiR-DNA dye, shown as a maximum intensity projection. Time in hours:mins, scale bar 10 µm. Black dashed circles show pronuclei, white denotes zygote outline. White arrows mark lagging chromosomes, yellow arrows mark polar bodies, orange arrows mark micronuclei and blue arrows denote nuclear defects. b) Incidence of no error vs. mitotic errors seen in the first mitotic division of embryos treated with DMSO or 2µM UMK57. P value from Fisher’s exact test. Confocal microscopy allows for increased detection of mitotic errors compared to previous widefield imaging. c) Types of mitotic errors seen in embryos treated with DMSO or 2µM UMK57. d) Maternal ages of zygotes in both treatment groups. P value from a Mann-Whitney U test. Green circles represent embryos with a chromosome segregation error occurring in mitosis 1, white circles showed no errors in mitosis 1. e) Metaphase plate width of control or UMK57 treated embryos at the time frame directly before anaphase, in embryos where metaphase plates were not rotated in 3D. Schematic provides visual representation of measurement taken (*d*). Pink circles represent embryos where a mitotic error occurred at the subsequent anaphase, white circles represent embryos with no mitotic errors. P value from a Mann-Whitney U test. f) Metaphase plate width of embryos which progressed through the first mitosis in the presence or absence of mitotic chromosome segregation errors. Pink circles represent 2µM UMK57 treated embryos, white circles represent DMSO treated (control) embryos.

Importantly, we observed significantly fewer chromosome mis-segregation events in the presence of UMK57 (Fig. 1a, bottom panel, 1b, n = 7 vs 19 embryos, p=0.002), and found a significant increase in the number of embryos progressing through the first division with no visible mitotic errors (Fig. 1b, 73% vs 29%, p=0.002). When subdividing by severity of chromosome error, we found that multipolar divisions and those with multiple lagging chromosomes were abolished in the UMK57-treated group (Fig. 1c).

This reduction in errors was not due to any change in the duration of the first mitosis (Fig. S1a) nor was it associated with differences in maternal age (Fig. 1d, S1b). Because previous work had implicated spindle pole defocusing in zygotic errors^25^, we measured metaphase plate diameter as a proxy of spindle geometry. We found a significant decrease upon treatment with UMK57 (Fig. 1e; p=0.003). However, increased metaphase plate width was not significantly associated with lagging chromosomes or multipolar chromosome segregation in our dataset (Fig. 1f; p=0.16). Instead, our data suggest that the hyper-stabilisation of kinetochore-microtubule attachments is more closely linked to the high frequency of these error-generating phenotypes in the first mitotic division than abnormal spindle geometry alone. Indeed, we suspect that UMK57 may also rescue abnormal spindle geometry but have not investigated this directly due to scarcity of human embryos.

**Figure S1.**
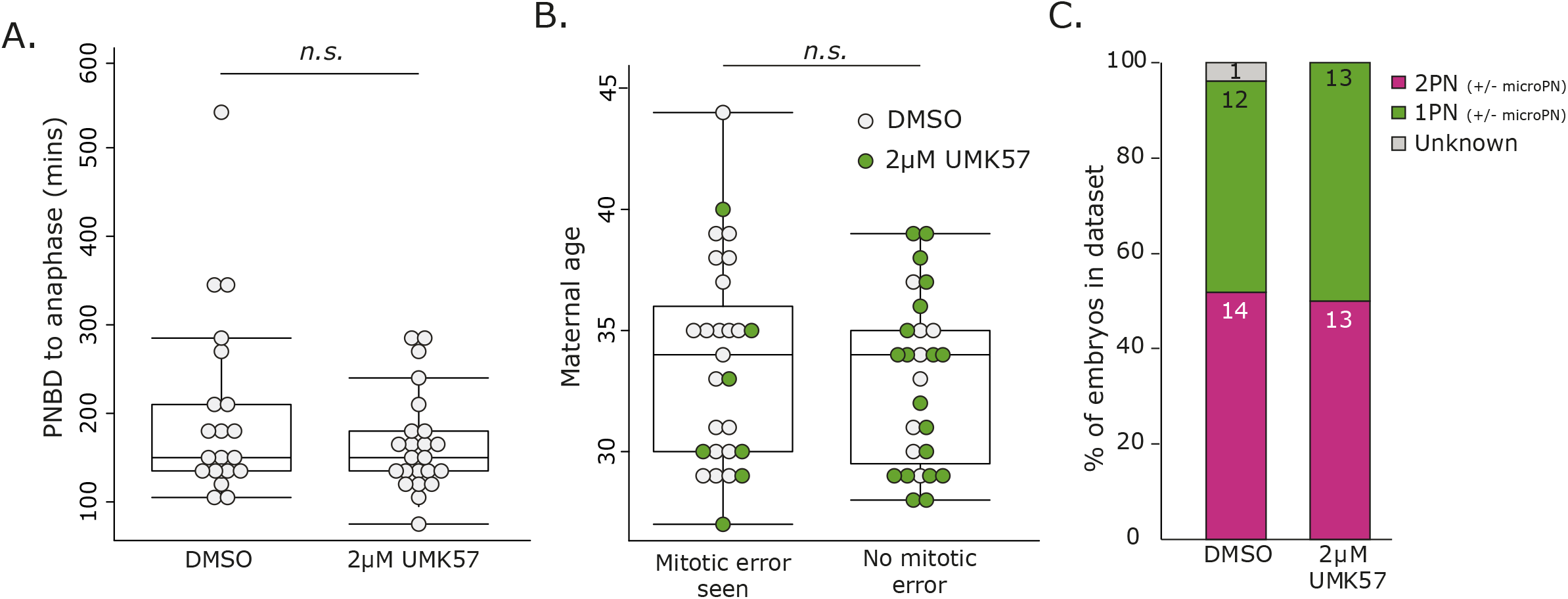
Supporting data. a) Mitotic timing from pronuclear breakdown (PNBD) to anaphase onset, as seen by live chromosome imaging in the first mitosis of human embryos. P value from a Mann-Whitney U test. b) Maternal ages of zygotes in groups where embryos either had a visible mitotic error in mitosis 1, or did not (as seen by live chromosome imaging). P value from a Mann-Whitney U test. Green circles represent embryos treated with 2µM UMK57, grey circles showed embryos treated with DMSO. c) Zygote pronuclei number in each treatment group. MicroPronuclei (microPN) are significantly smaller than the main pronuclei and contain a small amount of genetic material as visualised by SiR-DNA in live imaging. We assume these arise from chromosome errors in meiosis II and encapsulate single/low numbers of chromosomes.

### Aberrant cleavage patterns are ameliorated in UMK57 treated human embryos

Visual inspection of live imaging sequences of UMK57 treated human zygotes highlighted that all the divisions generated two daughter cells. This contrasts with the frequent observation of alternative cleavage patterns in the first division of the embryo in assisted reproduction cycles^37^. Examples of these from our clinical imaging are shown in Fig. 2a. We observed from research embryos cleaving abnormally that irregular cell formation is associated with chromosome segregation errors (Fig. 2b). However, our imaging revealed that apparently good quality 2-cell embryos can also be subject to chaotic mis-segregation (Fig. 2c). In the control group of embryos which divided into two cells (n=19), 40% had no mitotic errors, but 60% showed lagging chromosomes or multipolar segregation (Fig 2d). This demonstrates that chromosome errors and cell formation are not precisely coupled. As expected, 100% of abnormally cleaving embryos (n=8) displayed mitotic errors (Fig 2d).

**Figure 2.**
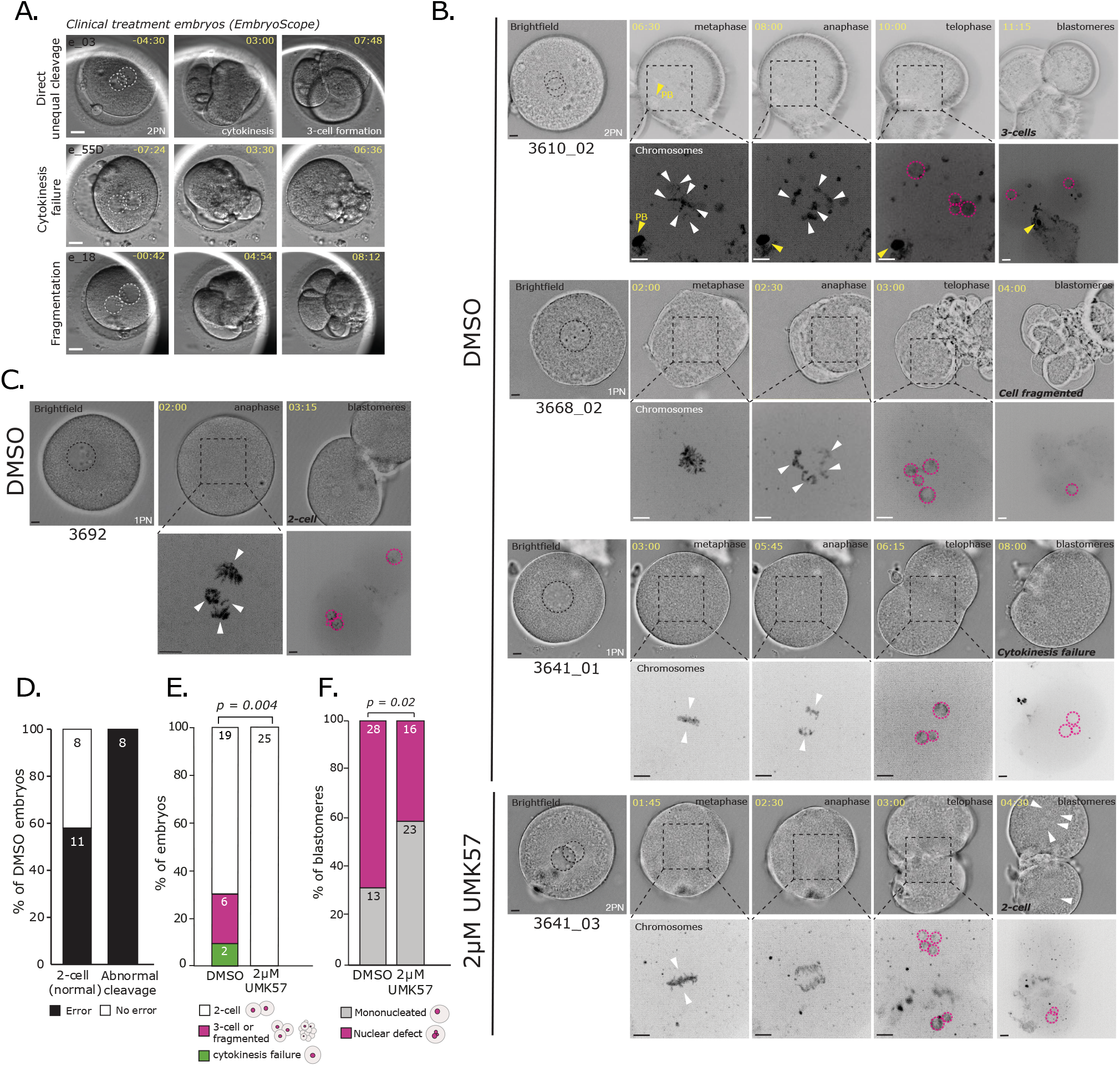
Rescue of chromosome errors results in an increased proportion of 2-cell embryos. a) Examples of abnormal first cleavage divisions in clinical 2PN embryos cultured via EmbryoScope for patient treatment in the IVF clinic. Time in hh:mm, scale bar 20µm. b) Representative images of research embryos progressing through the first mitosis with chromosomes labelled. DMSO treated embryos show similar phenotypes to abnormal treatment embryos. Time in hh:mm, scale bars all 10µm. Chromosome segregation errors are labelled with white arrows, nuclear defects at the 2-cell stage are labelled with pink dashed circles. Note nuclear defects occur in 2µM UMK57 treated embryo in the absence of chromosome errors. All brightfield images are a single Z slice, while chromosome images are Z projections c) Representative images of embryo 3692 progressing through the first cleavage in the presence of chaotic mitotic chromosome segregation (white arrows). Time in hh:mm, scale bar 10µm. Note two clear blastomeres are formed from this division. d) Mitotic chromosome errors in embryos showing normal or abnormal cell formation following the first mitosis. e) Embryo phenotypes following the first cleavage division in DMSO and 2µM UMK57 treated embryos. P-value from Fisher’s exact test (comparing frequency of 2-cell vs non 2-cell embryos per treatment group). f) Nuclear defects in DMSO and 2µM UMK57 treated embryos at the 2-cell stage. P value from a Fisher’s exact test.

Abnormally cleaving control zygotes divided directly into three cells, showed failed cytokinesis or had severe cytoplasmic fragmentation (Fig. 2e, n=8/19). Importantly, cell formation defects were eliminated upon UMK57 treatment, and 100% of zygotes in this group formed two cells following the first mitosis (Fig. 2e, n=25/25, p=0.004). The frequency of multinucleation in daughter blastomeres was also significantly reduced by UMK57 treatment (Fig. 2f, p=0.02). Again, this is consistent with a hyper-stable microtubule origin of these abnormal phenotypes, likely due to mitotic spindle dysfunction (see above and Ref.^25^). Why reducing microtubule stability would rescue cytokinesis failure is unclear. However the central spindle has been suggested to be important in defining cleavage planes^38^, and we speculate spindle dysfunction disrupts this process.

### Establishment of live cell imaging single cell sequencing to investigate karyotype evolution

To directly investigate the impact of error generating phenotypes on the karyotype of daughter blastomeres, we established a single-cell live imaging next generation sequencing assay using a commercial PGT-A platform (scLiveSeq). The schematic in Fig. 3a shows how zygotes were monitored through the first mitosis by live cell imaging and then triaged for sequencing. Embryos forming two distinct cells were disaggregated and each blastomere was sequenced separately to establish the karyotype. Abnormally cleaving embryos could not be readily disaggregated and are therefore not captured by this experiment.

**Figure 3.**
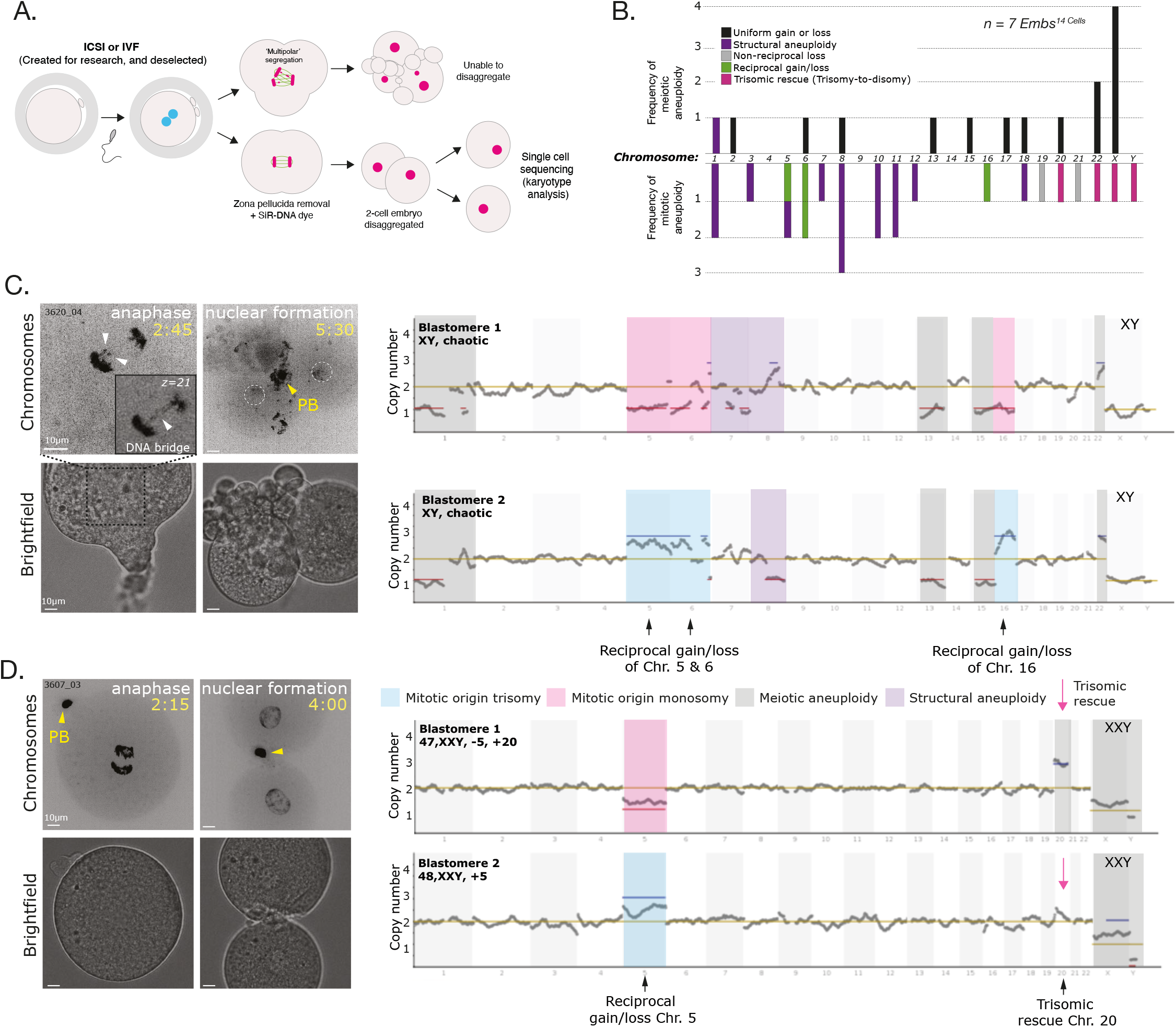
Karyotype evolution via mitotic-origin aneuploidy arising in the first human embryonic mitosis. a) Schematic of sc-LiveSeq assay. Human zygotes are prepared for live chromosome imaging by removing the zona pellucida and incubating with SiR-DNA dye, then monitored by confocal spinning disc microscopy through the first embryonic mitosis. Following the first divison, embryos are assessed for suitability for single cell sequencing. Embryos undergoing chaotic mitoses which result in fragmented and erroneous cell formation and are unable to be processed for sequencing. Embryos forming 2 clear cells are disaggregated for single cell sequencing. b) Frequency of mitotic- and meitoic-origin aneuploidies identified with next-generation sequencing in the cohort of DMSO treated embryos. c) Representative images from time lapse imaging of a human embryo (3620_04) progressing through the first embryonic mitosis with a complex mitotic error. Time in hh:mm, chromosomes are visualised using SiR-DNA dye, shown as a maximum intensity projection, scale bar 10 µm. Lagging chromosomes are marked with white arrows (Z projection), while inset image shows DNA bridge at z slice 21. Adjacent; single cell sequencing traces from the two blastomeres formed by embryo 3620_04. d) Representative images from time lapse imaging of a human embryo human embryo (3607_03) progressing through the first embryonic mitosis, with no mitotic errors. Time in hh:mm, chromosomes are visualised using SiR-DNA dye, shown as a maximum intensity projection, scale bar 10 µm. Yellow arrow denotes the polar body. Adjacent; single cell sequencing traces from the two blastomeres formed by embryo 3607_03. Trisomic rescue (pink arrow) describes a meiotic trisomy (Chr. 20) reverted to disomy in one resulting blastomere, via a mitotic-origin chromosome loss event. Note sex chromosome aneuploidy (XXY - elevated X, depressed Y signature) in this embryo.

We first determined the incidence of meiotic-origin aneuploidy in our dataset by identifying pairs of daughter blastomeres which both contained the same chromosome gain or loss (n=7 embryos, Fig. 3b). The fourteen detected numerical meiotic aneuploidies were found for both autosomes and sex chromosomes (Fig. 3b, black bars). We then determined to what extent mitotic-origin aneuploidy was found in the same embryos. We found ten cases where chromosome number was unbalanced at the 2-cell stage (Fig. 3b, green, grey and pink bars). This included four incidences of reciprocal gain/loss (green bars) which give rise to trisomy and monosomy, respectively. We also identified four cases of meiotic trisomic rescue at the 2-cell stage (pink bars). This is an event where loss of a chromosome ‘corrects’ a trisomy generated during meiosis, converting it into a disomy. We observed trisomic rescue of small autosomes (Chr 20 and 22) and sex chromosomes. For example, in embryo 3684_03, there was loss of a Y chromosome during the first division of the embryo creating a mosaic XXY/XX (Klinefelter) 2-cell embryo. To our knowledge, this is the first direct evidence of trisomic rescue occurring at the first mitotic division of the human embryo; indirect evidence comes from lineage tracing of somatic mutations from human foetal and placental samples^39^. We also identified thirteen mitotic-origin structural aneuploidies in our dataset of 2-cell embryos (purple bars), consistent with increased genomic instability at the zygote stage^26^ (Fig. 3b). These were predominantly partial chromosome losses, but we saw evidence for partial chromosome gains and translocations. These combined data provide direct evidence of numerical and structural mosaic aneuploidy arising from the first mitotic division of human embryos.

Because we had live chromosome imaging of anaphase for some of these embryos, we could directly assess the relationship between mitotic phenotypes and karyotype change. We observed chaotic mitotic aneuploidy following an error-prone division, confirming that lagging chromosomes do cause aneuploidy (Fig. 3c). This embryo also had an anaphase DNA bridge which was likely associated with the multiple structural aneuploidies seen.

Surprisingly, we could also detect karyotype change when there was no observable error-generating phenotype (Fig. 3d). This likely reflects mis-segregating chromosomes being hidden within the anaphase masses, an event also reported in somatic cells^40^.

### Restored mitotic fidelity decreases mosaic aneuploidy in 2-cell human embryos

Considering the significant decrease in mitotic error phenotypes following treatment with UMK57, we used sc-LiveSeq to ask whether there is also a change in the incidence of mitotic-origin aneuploidy. In the presence of UMK57, we still detected eight cases of meiotic-origin aneuploidies, consistent with control embryos (Fig. 4a,b, black bars). However, we could observe divisions where a lagging chromosome was seemingly corrected in anaphase and did not result in mosaic aneuploidy, suggesting restoration of functional error-correction pathways (Fig. 4a). We then asked what fraction of 2-cell embryos contained a blastomere with no mitotic origin aneuploidies. Treatment with UMK57 resulted in 6/12 embryos with no errors, compared to 1/14 in the control (Fig. 4c, p=0.02). Thus, UMK57 can attenuate mitotic origin aneuploidy.

**Figure 4.**
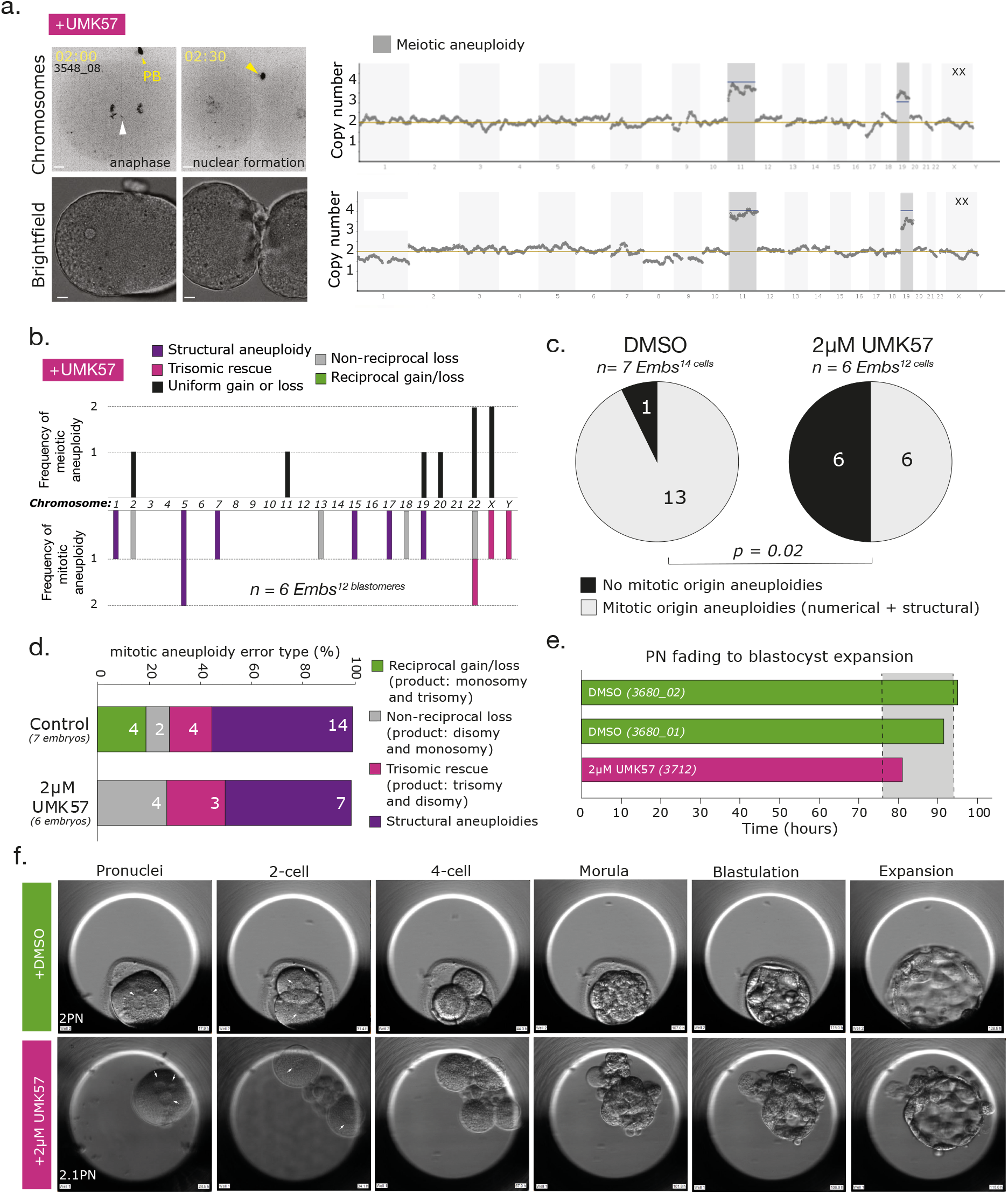
Rescue of mosaic aneuploidy in 2-cell human embryos. a) sc-LiveSeq representative images from 2µM UMK57 treated embryo (3548_08) undergoing the first embryonic mitosis, with a lagging chromosome (white arrow). Polar body marked by yellow arrow. Scale bar 10µm. Adjacent: representative single cell sequencing traces for karyotype from blastomeres resulting from embryo 3548_08, showing an absence of mitotic-origin aneuploidies. b) Frequency of mitotic- and meiotic-origin aneuploidies in the cohort of 2µM UMK57 treated embryos. c) Overall frequency of blastomeres displaying mitotic-origin aneuploidies (defined as whole chromosome (numerical) plus structural aneuploidies) in the cohorts of DMSO and 2µM UMK57 treated 2-cell embryos. P value from a binomial test. d) Frequency of different mitotic-origin aneuploidies in DMSO and 2µM UMK57 treated embryos. Reciprocal gain/loss is defined as reciprocal monosomy/trisomy of a chromosome in a single embryo. Non-reciprocal loss is defined as disomy/monosomy of a chromosome in a single embryo. Trisomic rescue is rescue of a meiotic trisomy to disomy through non-reciprocal chromosome loss. e) Timings of blastocyst formation in DMSO and 2µM UMK57, as observed using EmbryoScope time lapse imaging. Grey box shows average time ascertained from embryoscope imaging of clinical treatment embryos (Desai *et al*, 2014 PMID: 24951056). f) Representative images of deselected zygotes progressing to blastocyst stage in the presence of DMSO or 2µM UMK57, observed with EmbryoScope. Zona pellucida was removed from UMK57 treated embryo to ensure permeation of the drug.

Inspection of the error types indicated that non-reciprocal loss (grey bars) and trisomic rescue (pink bars) were present in both groups, suggesting these processes are not rescued by UMK57 (Fig. 4b, d). However, the number of structural aneuploidies in UMK57-treated embryos was halved compared to the control group (purple bars, 7 vs 13, Fig. 3b, 4b, d).

Moreover, reciprocal gain/loss (green bars) were not observed in the cohort of 2µM UMK57 treated embryos (Fig. 4b, d); therefore, UMK57 significantly reduced the true non-disjunction events (p=0.045). While underpowered, these data hint at a reduction in error rate and a change in chromosome error type when MCAK activity is increased. Aneuploidies were in fact more likely to be clustered within a single daughter cell originating from a mother zygote (p<0.05, pooling cells from both treatment groups). *i*.*e*. if a blastomere has a numerical aneuploidy error it is more likely to have others of same type (on different chromosomes). In fact observing as many as 6 (50%) blastomeres with no numerical or structural aneuploidies in UMK57 (Fig 4c) is dependent on error clustering (p<0.05). This suggests the first mitotic division is unequal at distributing errors between daughter cells and might be indicative of spindle asymmetry.

As previously mentioned, our single cell sequencing assay is only compatible with embryos that form two distinct blastomeres, thus selecting against chaotic divisions in control embryos which, importantly, were not observed in UMK57 treated embryos (Fig. 2d, 3a).

Therefore, the true reduction in mitotic aneuploidy is likely to be higher. Finally, we cultured a human zygote to the blastocyst stage in the presence of 2µM UMK57, demonstrating compatibility with successful preimplantation embryo development (Fig. 4e, f).

## Discussion and conclusions

Here we combined live cell chromosome imaging and single cell sequencing of human 2-cell embryos to reveal how errors in the first mitotic division cause karyotypic change. This confirms that the first mitosis is a key driver of mosaic aneuploidy – a feature of >80% human preimplantation embryos^41^. We were able to detect both structural rearrangements and whole chromosome losses and/or gains (generating monosomic and trisomic lineages in the embryo). Our experiments also show that chromosome loss events can “rescue” a trisomy that occurred during the meiotic divisions. The result is a 2-cell embryo which now contains both a trisomic and diploid lineage for the affected chromosome. While we cannot detect uniparental disomy incidence with our assay, this raises the possibility that mitotic errors can have a positive effect by removing trisomies that would otherwise lead to human developmental disorders or lethality.

While low/mid-level mosaic blastocysts can give rise to healthy live birth^17^, chaotic mitotic-origin aneuploidy is associated with early embryo arrest^19^. Moreover, the developmental potential of blastocysts is negatively impacted by alternative cleavage patterns during the first divisions^42^. Clinical embryos showing severe problems in the first cleavage have an 8% chance of forming a blastocyst and 0% chance of giving rise to live birth^31^.We demonstrate that the small molecule agonist UMK57 can prevent these detrimental phenotypes. All UMK57-treated zygotes gave rise to 2-cell embryos with reduced multi-nucleation and an increased probability of containing a euploid cell lineage (Fig. 5). This advance will allow for controlled experimentation to uncover the role of aneuploidy in early bottlenecks of human reproduction (including implantation) and provides a potential intervention to improve the efficiency of blastocyst formation in IVF.

**Figure 5.**
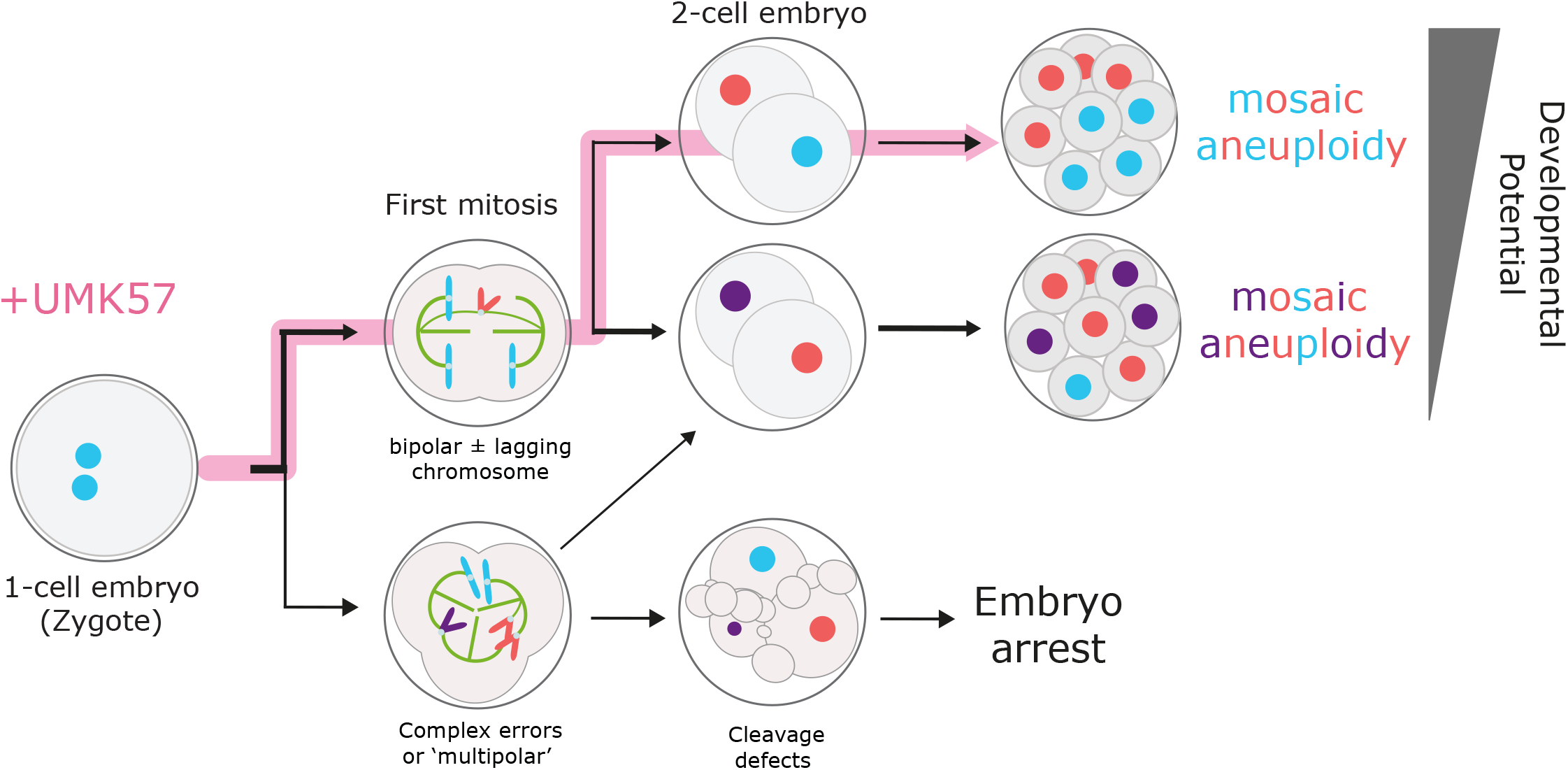
Model of how UMK57 can shift the developmental potential of human embryos. Human zygotes (2PN) undergo the first mitotic division leading to either bipolar divisions (green background) with or without lagging chromosomes (coloured red) (top; the number of which can vary in severity) or undergo multipolar chromosome segregation or segregations into multiple nuclei (bottom). The frequency is ~75% vs ~25%, respectively (indicated by stroke of the black arrows). Bipolar divisions (top) can give rise to 2-cell embryos where blastomeres are either both aneuploid (lower; red and purple nuclei; ~90% of the time) or contain an aneuploid (numerical or structural) and euploid lineage (upper; blue is euploid nuclei; ~10% of the time). These give rise to embryos with different severities of mosaicism which relates to the developmental potential and future pregnancy outcome. Multipolar divisions (bottom - blue background) give rise to either 2-cell embryos with incorrect nuclei numbers or embryos with defective cleavage patterns (direct cleavage to three cells, cytokinesis failure or severe cytoplasmic blebbing). The latter are likely to arrest before the formation of a blastocyst and are also likely to be aneuploid. Treatment of human zygotes with UMK57 biases the developmental pathway as indicated by the yellow arrow. All zygotes execute a bipolar division into 2-cells and the number with a euploid chromosome complement is now 50%. This is expected to increase the number of human embryos with high developmental potential in patients undergoing IVF treatment for infertility.

UMK57 activates the microtubule depolymerase KIF2C which functions to increase the turnover of microtubule-kinetochore attachments during mitosis. This improves the efficiency of error correction and reduces chromosome mis-segregation in cancer, stem and aged cells^28,32,33^. It follows that, 1) human zygotes also have hyperstable kinetochore microtubule attachments and this explains the high rate of chromosome mis-segregation and 2) human zygotes must possess the necessary biological machinery to correctly segregate chromosomes. Consistent with this, a patient identified as having a gain-of-function mutation in Aurora B (a key error-correction kinase) presented with ‘lower than average’ levels of aneuploidy in their IVF treatment embryos^43^. UMK57 also reduced the burden of structural aneuploidy, which is generally associated with replication stress and DNA damage^26^. One possibility is that the breakpoints reflect known fragile sites - common sites of DNA damage in human embryos^44^. It follows that a more dynamic spindle may reduce the risk of chromosome breakage during mitosis, consistent with experimental work in mouse^45^. The additional reversal of cell division defects and multinucleation in zygotes suggests that UMK57 treatment improves multiple aspects of mitotic spindle function.

In conclusion, we have shown that deregulation of the microtubule cytoskeleton is a key driver for generation of mosaic aneuploidy and abnormal embryo cleavage patterns. Future work will establish whether UMK57 treatment can override cell division and aneuploidy-associated embryo arrest, which is of great interest to the assisted reproduction field.

## Limitations

Access to human embryos for research is extremely challenging thus leading to limitations in the number of replicates. We have used both embryos created for research purposes via ICSI, and deselected embryos where fertilisation was delayed or deviated in number of pronuclei (e.g. 1PNs and/or zygotes containing micro-pronuclei) (Fig. S1c). Importantly, we excluded embryos where the number of major pronuclei was above two (e.g. 3PNs), as polyspermy and extra-numerary centrosomes are suspected here. We also excluded embryos where cytokinesis was premature and not linked to mitotic events. Our sequencing dataset is biased towards embryos where there is accurate cell division into two blastomeres that can be disaggregated. 1 in 4 of the control embryos (and none in UMK57 treated) undergo catastrophic first cleavage, are almost certainly aneuploid and are not captured in our sequencing dataset. Thus, the magnitude of the error-correcting effect of UMK57 based on single cell sequencing is likely to be an underestimate.

## METHODS

### Donation of human embryos to research

The NHS Research Ethics Committee approved both the research project (04/Q2802/26) and egg sharing to research programme (19/WM/0003). All work was conducted under a Research Licence from the Human Fertilisation and Embryology Authority (HFEA; R0155). Informed consent for donation of eggs, embryos and sperm to research was provided by patients undergoing in vitro fertilisation (IVF) or intracytoplasmic sperm injection (ICSI) at the Centre for Reproductive Medicine (CRM), University Hospitals Coventry and Warwickshire (UHCW) NHS Trust. Details of the CRM clinic success rates are published by the Human Fertilisation and Embryology Authority (https://www.hfea.gov.uk/choose-a-clinic/clinic-search/results/13/). Standard clinical protocols were used, including ovarian stimulation with FSH preparations according to either GnRH agonist or antagonist regimens, and the Origio (Cooper Surgical) or Gx (Vitrolife) suite of culture media. The deselected material used for research was unsuitable for the patients’ treatment, due to delay (0PN) or mis-fertilisation (1PN/microPN), and would otherwise have been disposed of. 1PN embryos may be haploid, but if diploid, may arise from fusion of the parental pronuclei and can lead to live birth^46^. Egg-share material was collected from volunteer patients aged ≤33 on the egg sharing to research programme at the CRM only, as described previously^3^. The majority of donated material was used fresh, but some unfertilised mature eggs were cryopreserved and later thawed, using Kitazato vitrification (VT601) and thawing (VT602) kits. Furthermore, 17 cryopreserved mature oocytes were altruistically donated to this project by a single patient. For fresh and thawed donated oocytes, mature (metaphase II) eggs were inseminated with fertile donor sperm via ICSI using standard clinical protocols in order to create embryos for research use in this project. One sperm donor was used for all research inseminations. Following sperm injection, eggs were incubated in a EmbryoScope™ time lapse incubator in an identical manner to embryos used clinically. Immature eggs collected at the same time from egg sharers were not used for research included in this publication.

### Live chromosome imaging of human embryos

Embryos arising from the egg share programme, or donated frozen oocytes, were collected from the clinic 1-2 hours after the fertilisation check had been completed by a clinical embryologist ~17 hours after insemination. Deselected embryos were collected from the clinic around 3-5 hours after the fertilisation check. For SiR-DNA treatment, the zona pellucida of individual embryos was removed by brief incubation and pipetting in pre-warmed acid Tyrode’s solution (Sigma). Embryos were transferred to a Fluorodish (WPI) containing 2 µM SiR-DNA (Spirochrome) diluted in pre-warmed Cleav medium (Origio) or GxTL (Vitrolife) under mineral oil. Embryos were transported ~14 km from UHCW CRM to Warwick Medical School (WMS) in a portable incubator (K Systems) held at 37ºC, and transferred to the microscope immediately upon arrival. Embryos were imaged using Marianas spinning disc confocal microscope from 3i (Intelligent Imaging Innovations) equipped with 2x Photometrics 95B Prime sCMOS cameras, using a 40x oil-immersion 1.46, alphaPlnApo (Zeiss) objective. Fluorescent images were acquired every 15 minutes (60 × 1.5 µm optical sections) with a 640 nm laser attenuated to 4% with an exposure time of 40 ms, and brightfield images were acquired attenuated to 4% with an exposure time of 40 ms. A stage-top incubator (Okolab) maintained embryos at 37ºC and 5% CO_2_. Temperature and CO_2_ were verified using calibrated probes. Image sequences were analysed by hand.

### Drug treatments

UMK57 (342595-74-8, MedKoo Biosciences Inc.) was diluted in DMSO (D2438, Sigma Aldrich) to a working stock concentration of 2 mM. The final experimental concentration was 2 µM, diluted in Origo or GX-TL IVF culture media. The same volume of DMSO was used as a control. Embryos were incubated with UMK57 or DMSO at either the MII or PN stage, depending on maturity at time of sample collection, following removal of the zona pellucida as described above.

### Embryoscope culture and imaging

Culture media (Gx-TL) containing either DMSO or 2 µM UMK57 was used to equilibrate a 12-well EmbryoScope® time-lapse culture dish overlaid with mineral oil. Zygotes were prepared for treatment with either DMSO or 2 µM UMK57 as above, moved to the EmbryoScope® dish, and transported to Warwick Medical School as described above.

Zygotes were transferred to the research EmbryoScope immediately upon arrival and cultured for 5-6 days with no media changes, continuously monitored by 10 min brightfield time-lapse imaging. Time-lapse sequences were analysed by hand.

### 2-cell human embryo ploidy analysis

Following live imaging (either confocal or EmbryoScope), embryos which formed two blastomeres without extensive fragmentation were selected for sequencing. Disaggregation and tubing were performed in a Class 2 microbiology safety cabinet following decontamination using DNAse-Away (7010, Thermo Fisher) and UV. 2-cell embryos were placed in a 7 µL drop of wash buffer (1x PBS/0.1% BSA), and disaggregated by pipetting using a 135 µm needle and Stripper pipette (CooperSurgical). Blastomeres were individually washed and transferred to a sterile Eppendorf PCR tube. A sample of the final wash drop was tubed separately to control for sterility. Samples were held at −20°C and shipped to the Igenomix laboratory (Guildford UK, or Valencia, Spain) on ice.

PGT-A analysis for single cell samples was performed by Igenomix (Guildford, UK or Valenica, Spain) - a clinical testing laboratory - using a next generation sequencing (NGS) platform. The NGS platform used to analyse samples in this study was a semiautomated protocol using the Ion Chef™ equipment for library preparation and the S5 sequencer (ThermoFisher Scientific, Waltham, MA, USA). Samples were tested in batches of 24 or 96 (520 and 530 chips, respectively). Whole genome amplification (WGA) and DNA barcoding were performed using the Ion ReproSeq PGS Kit (ThermoFisher Scientific), following the manufacturer’s instructions. The amplified DNA was purified, quantified with the Qubit™ (Qubit dsDNA HS Assay Kit, ThermoFisher Scientific), and diluted before placing it in the Ion Chef™ instrument that automates preparation of the library and templates, as well as chip loading. The sequencing run and analysis process were automatically completed following the ReproSeq PGS w1.1 workflow by the Torrent Server and the Ion Reporter in Torrent Suite™ Software (Life Technologies Ltd., Paisley, United Kingdom). The final output result for each sample was a copy number diagram for all chromosomes.

### Data and Statistical Analysis

The Mann-Whitney U tests for Fig 1d, 1e, Fisher’s exact tests for Fig. 1b, 2e, 2f and 4d and binomial test for Fig 4c, were performed using MATLAB R2020A (Mathworks) inbuilt functions. Metaphase plate width in Fig. 1e was measured at the time point immediately before anaphase onset, using the line draw tool in SlideBook 2023 (3i Intelligent Imaging Innovations). Clustering of errors was analysed using a goodness-of-fit test on cells with errors, sampling the null hypothesis that errors are independent. A Monte Carlo simulation randomly distributing the 4 error types (reciprocal gain/loss, single losses, paired structural aneuploidy, unpaired structural aneuploidy) was used to analyse the probability of observing a given number of cells that were free of aneuploidies in DMSO and UMK57 groups. All plots were created in Excel or MATLAB, and figures were created using Adobe Illustrator.

## CRediT Author Contributions

**Cerys Currie:** Conceptualisation, Methodology, Investigation, Data collection and curation, Validation, Formal Analysis, Original Draft Preparation, Writing and Editing, **Aleksandra Byrska:** Methodology (preparation of samples for sequencing), Data Collection, Data Validation **Debbie Taylor:** Methodology, Resources, **Muriel Erent:** Project administration, **Daniela Bakalova and Xuhui Sun (Igenomix):** Service provision of clinical PGT-A, **Constandina Koki:** Statistics support and analysis, **Nigel Burroughs:** Statistics support and analysis **Richard Anderson:** Reviewing and Editing, Funding acquisition, **Adele Marston**: Reviewing and Editing, Funding acquisition, **Geraldine Hartshorne:** Reviewing and Editing, Supervision, Ethical approvals, Funding acquisition, **Andrew McAinsh:** Conceptualisation, Supervision, Original Draft Preparation, Writing and Editing, Funding acquisition.

## ACKNOWLEDGEMENTS

We are hugely thankful and indebted to the patients, embryologists, research nurses and medical consultants at the Centre for Reproductive Medicine, University Hospitals Coventry and Warwickshire NHS Trust. Special thanks to all Warwick and Edinburgh colleagues in the Eggs ‘n Embryos research group for insightful and fun discussions – supported by a Wellcome Collaborative Award (215625). We also gratefully acknowledge CAMDU (Computing and Advanced Microscopy Unit) for their support and assistance in this work, particularly Maëlle Lorvellec for microscopy assistance and Laura Cooper for data assistance. We thank Thomas Keen for laboratory assistance. A.D.M, A.L.M., G.M.H., R.A.A., C.E.C., and D.T. are supported by a Wellcome Collaborative Award (215625). A.B. is supported by a Warwick Chancellor’s International Scholarship. G.M.H. is also supported by the WPH Charitable Trust. A.L.M. is also supported by a Wellcome Investigator award (220780) and funding for the Discovery Research Platform for Hidden Cell Biology (226791).

---

**Table 1.**
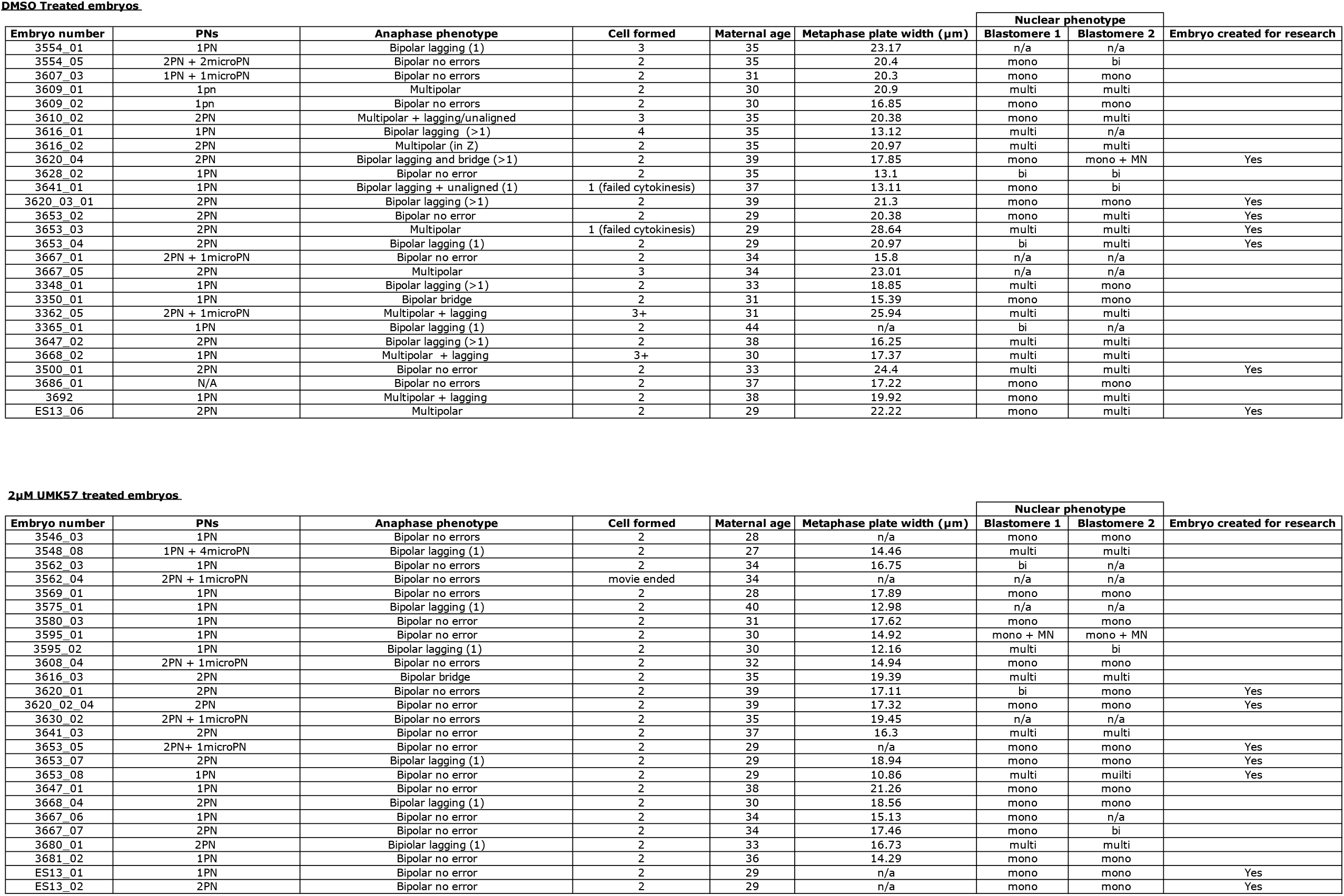
Chromosome imaging analysis of human zygotes treated with either DMSO (top) or 2µM UMK57 (bottom).

**DMSO Treated embryos.**
| Embryo number | PNs | Anaphase phenotype | Cell formed | Maternal age | Metaphase plate width (μm) | Nuclear phenotype |  | Embryo created for research |
| --- | --- | --- | --- | --- | --- | --- | --- | --- |
|  |  |  |  |  |  | Blastomere 1 | Blastomere 2 |  |
| 3554_01 | 1PN | Bipolar lagging (1) | 3 | 35 | 23.17 | n/a | n/a |  |
| 3554_05 | 2PN + 2microPN | Bipolar no errors | 2 | 35 | 20.4 | mono | bi |  |
| 3607_03 | 1PN + 1microPN | Bipolar no errors | 2 | 31 | 20.3 | mono | mono |  |
| 3609_01 | 1pn | Multipolar | 2 | 30 | 20.9 | multi | multi |  |
| 3609_02 | 1pn | Bipolar no errors | 2 | 30 | 16.85 | mono | mono |  |
| 3610_02 | 2PN | Multipolar + lagging/unaligned | 3 | 35 | 20.38 | mono | multi |  |
| 3616_01 | 1PN | Bipolar lagging (>1) | 4 | 35 | 13.12 | multi | n/a |  |
| 3616_02 | 2PN | Multipolar (in Z) | 2 | 35 | 20.97 | multi | multi |  |
| 3620_04 | 2PN | Bipolar lagging and bridge (>1) | 2 | 39 | 17.85 | mono | mono + MN | Yes |
| 3628_02 | 1PN | Bipolar no error | 2 | 35 | 13.1 | bi | bi |  |
| 3641_01 | 1PN | Bipolar lagging + unaligned (1) | 1 (failed cytokinesis) | 37 | 13.11 | mono | bi |  |
| 3620_03_01 | 2PN | Bipolar lagging (>1) | 2 | 39 | 21.3 | mono | mono | Yes |
| 3653_02 | 2PN | Bipolar no error | 2 | 29 | 20.38 | mono | multi | Yes |
| 3653_03 | 2PN | Multipolar | 1 (failed cytokinesis) | 29 | 28.64 | multi | multi | Yes |
| 3653_04 | 2PN | Bipolar lagging (1) | 2 | 29 | 20.97 | bi | multi | Yes |
| 3667_01 | 2PN + 1microPN | Bipolar no error | 2 | 34 | 15.8 | n/a | n/a |  |
| 3667_05 | 2PN | Multipolar | 3 | 34 | 23.01 | n/a | n/a |  |
| 3348_01 | 1PN | Bipolar lagging (>1) | 2 | 33 | 18.85 | multi | mono |  |
| 3350_01 | 1PN | Bipolar bridge | 2 | 31 | 15.39 | mono | mono |  |
| 3362_05 | 2PN + 1microPN | Multipolar + lagging | 3+ | 31 | 25.94 | multi | multi |  |
| 3365_01 | 1PN | Bipolar lagging (1) | 2 | 44 | n/a | bi | n/a |  |
| 3647_02 | 2PN | Bipolar lagging (>1) | 2 | 38 | 16.25 | multi | multi |  |
| 3668_02 | 1PN | Multipolar + lagging | 3+ | 30 | 17.37 | multi | multi |  |
| 3500_01 | 2PN | Bipolar no error | 2 | 33 | 24.4 | multi | multi | Yes |
| 3686_01 | N/A | Bipolar no errors | 2 | 37 | 17.22 | mono | mono |  |
| 3692 | 1PN | Multipolar + lagging | 2 | 38 | 19.92 | mono | multi |  |
| ES13_06 | 2PN | Multipolar | 2 | 29 | 22.22 | mono | multi | Yes |

| Embryo number | PNs | Anaphase phenotype | Cell formed | Maternal age | Metaphase plate width (μm) | Nuclear phenotype |  | Embryo created for research |
| --- | --- | --- | --- | --- | --- | --- | --- | --- |
|  |  |  |  |  |  | Blastomere 1 | Blastomere 2 |  |
| 3546_03 | 1PN | Bipolar no errors | 2 | 28 | n/a | mono | mono |  |
| 3548_08 | 1PN + 4microPN | Bipolar lagging (1) | 2 | 27 | 14.46 | multi | multi |  |
| 3562_03 | 1PN | Bipolar no errors | 2 | 34 | 16.75 | bi | n/a |  |
| 3562_04 | 2PN + 1microPN | Bipolar no errors | movie ended | 34 | n/a | n/a | n/a |  |
| 3569_01 | 1PN | Bipolar no errors | 2 | 28 | 17.89 | mono | mono |  |
| 3575_01 | 1PN | Bipolar lagging (1) | 2 | 40 | 12.98 | n/a | n/a |  |
| 3580_03 | 1PN | Bipolar no error | 2 | 31 | 17.62 | mono | mono |  |
| 3595_01 | 1PN | Bipolar no error | 2 | 30 | 14.92 | mono + MN | mono + MN |  |
| 3595_02 | 1PN | Bipolar lagging (1) | 2 | 30 | 12.16 | multi | bi |  |
| 3608_04 | 2PN + 1microPN | Bipolar no errors | 2 | 32 | 14.94 | mono | mono |  |
| 3616_03 | 2PN | Bipolar bridge | 2 | 35 | 19.39 | multi | multi |  |
| 3620_01 | 2PN | Bipolar no errors | 2 | 39 | 17.11 | bi | mono | Yes |
| 3620_02_04 | 2PN | Bipolar no error | 2 | 39 | 17.32 | mono | mono | Yes |
| 3630_02 | 2PN + 1microPN | Bipolar no errors | 2 | 35 | 19.45 | n/a | n/a |  |
| 3641_03 | 2PN | Bipolar no error | 2 | 37 | 16.3 | multi | multi |  |
| 3653_05 | 2PN+ 1microPN | Bipolar no error | 2 | 29 | n/a | mono | mono | Yes |
| 3653_07 | 2PN | Bipolar lagging (1) | 2 | 29 | 18.94 | mono | mono | Yes |
| 3653_08 | 1PN | Bipolar no error | 2 | 29 | 10.86 | multi | multi | Yes |
| 3647_01 | 1PN | Bipolar no error | 2 | 38 | 21.26 | mono | mono |  |
| 3668_04 | 2PN | Bipolar lagging (1) | 2 | 30 | 18.56 | mono | mono |  |
| 3667_06 | 1PN | Bipolar no error | 2 | 34 | 15.13 | mono | n/a |  |
| 3667_07 | 2PN | Bipolar no error | 2 | 34 | 17.46 | mono | bi |  |
| 3680_01 | 2PN | Bipolar lagging (1) | 2 | 33 | 16.73 | multi | multi |  |
| 3681_02 | 1PN | Bipolar no error | 2 | 36 | 14.29 | mono | mono |  |
| ES13_01 | 1PN | Bipolar no error | 2 | 29 | n/a | mono | mono | Yes |
| ES13_02 | 2PN | Bipolar no error | 2 | 29 | n/a | mono | mono | Yes |

**Table 2.**
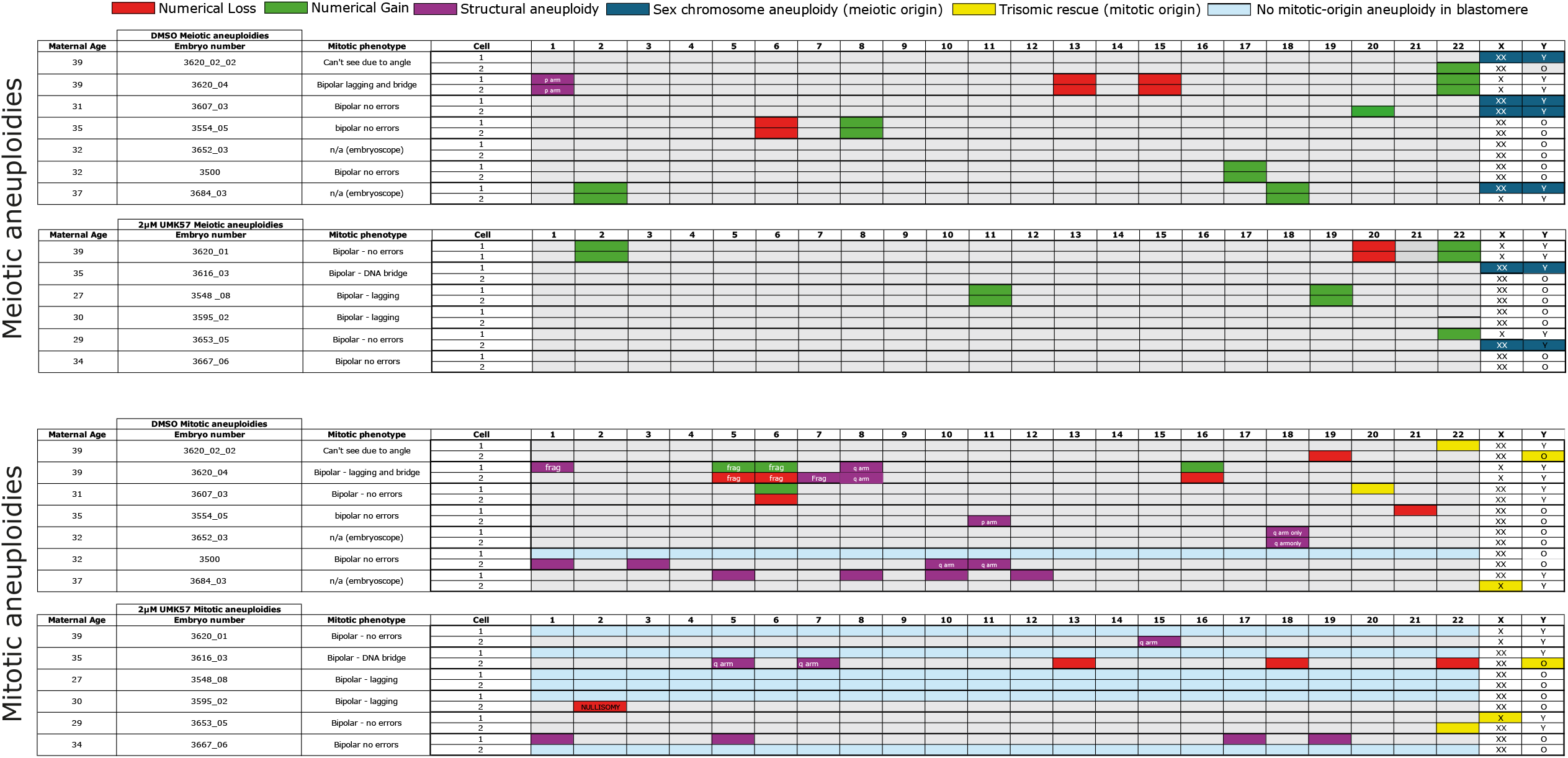
Summary table of meiotic and mitotic origin aneuploidies in human embryos treated with DMSO (top), or 2µM UMK57 (bottom). Blastomeres were disaggregated at the 2-cell stage and sent for karyoptype sequencing separately. Meiotic aneuploidies were identified as equally affecting both cells, or where a trisomy existed but was reverted to disomy in the corresponding daughter cell (trisomic rescue), as a single chromatid cannot be gained *de-novo*. Mitotic origin aneuploidies differed between daughter blastomeres.

